# A Histomorphometric and Computational Investigation of the Stabilizing Role of Pectinate Ligaments in the Aqueous Outflow Pathway

**DOI:** 10.1101/2023.10.17.562754

**Authors:** Babak N. Safa, Nina Sara Fraticelli Guzmán, Guorong Li, W. Daniel Stamer, Andrew J. Feola, C. Ross Ethier

## Abstract

Murine models are commonly used to study glaucoma, the leading cause of irreversible blindness. Glaucoma is associated with elevated intraocular pressure (IOP), which is regulated by the tissues of the aqueous outflow pathway. In particular, pectinate ligaments (PLs) connect the iris and trabecular meshwork (TM) at the anterior chamber angle, with an unknown role in maintenance of the biomechanical stability of the aqueous outflow pathway, thus motivating this study. We conducted histomorphometric analysis and optical coherence tomography-based finite element (FE) modeling on three cohorts of C57BL/6 mice: ‘young’ (2-6 months), ‘middle-aged’ (11-16 months), and ‘elderly’ (25-32 months). We evaluated the age-specific morphology of the outflow pathway tissues. Further, because of the known pressure-dependent Schlemm’s canal (SC) narrowing, we assessed the dependence of the SC lumen area to varying IOPs in age-specific FE models over a physiological range of TM/PL stiffness values. We found age-dependent changes in morphology of outflow tissues; notably, the PLs were more developed in older mice compared to younger ones. In addition, FE modeling demonstrated that murine SC patency is highly dependent on the presence of PLs, and that increased IOP caused SC collapse only with sufficiently low TM/PL stiffness values. Moreover, the elderly model showed more susceptibility to SC collapse compared to the younger models. In conclusion, our study elucidated the previously unexplored role of PLs in the aqueous outflow pathway, indicating their function in supporting TM and SC under elevated IOP.

## Introduction

Glaucoma is a group of eye diseases that are collectively responsible for the majority of irreversible blindness worldwide [1]. Glaucoma is often associated with increased intraocular pressure (IOP), i.e., ocular hypertension [2,3], and lowering of IOP helps preserve vision [4–7]. Therefore, understanding the mechanisms by which IOP is regulated has great importance in developing new glaucoma treatments.

A critical determinant of IOP is the hydrodynamic resistance of the conventional outflow pathway tissues, including the inner wall of Schlemm’s canal (SC) and trabecular meshwork (TM; Figure 1). Further, the biomechanical properties of these tissues are thought to play a key role in IOP homeostasis [8,9]. For example, several studies have observed TM stiffening in glaucoma and mechanobiological factors are known to affect aqueous humor drainage across the inner wall of SC [10–13]. Another integral part of the outflow pathway in many species is the pectinate ligaments (also known as iris processes [14]), yet little is known about their biomechanical properties and their role in ocular hypertension and the regulation of IOP [14–19].

**Figure 1:**
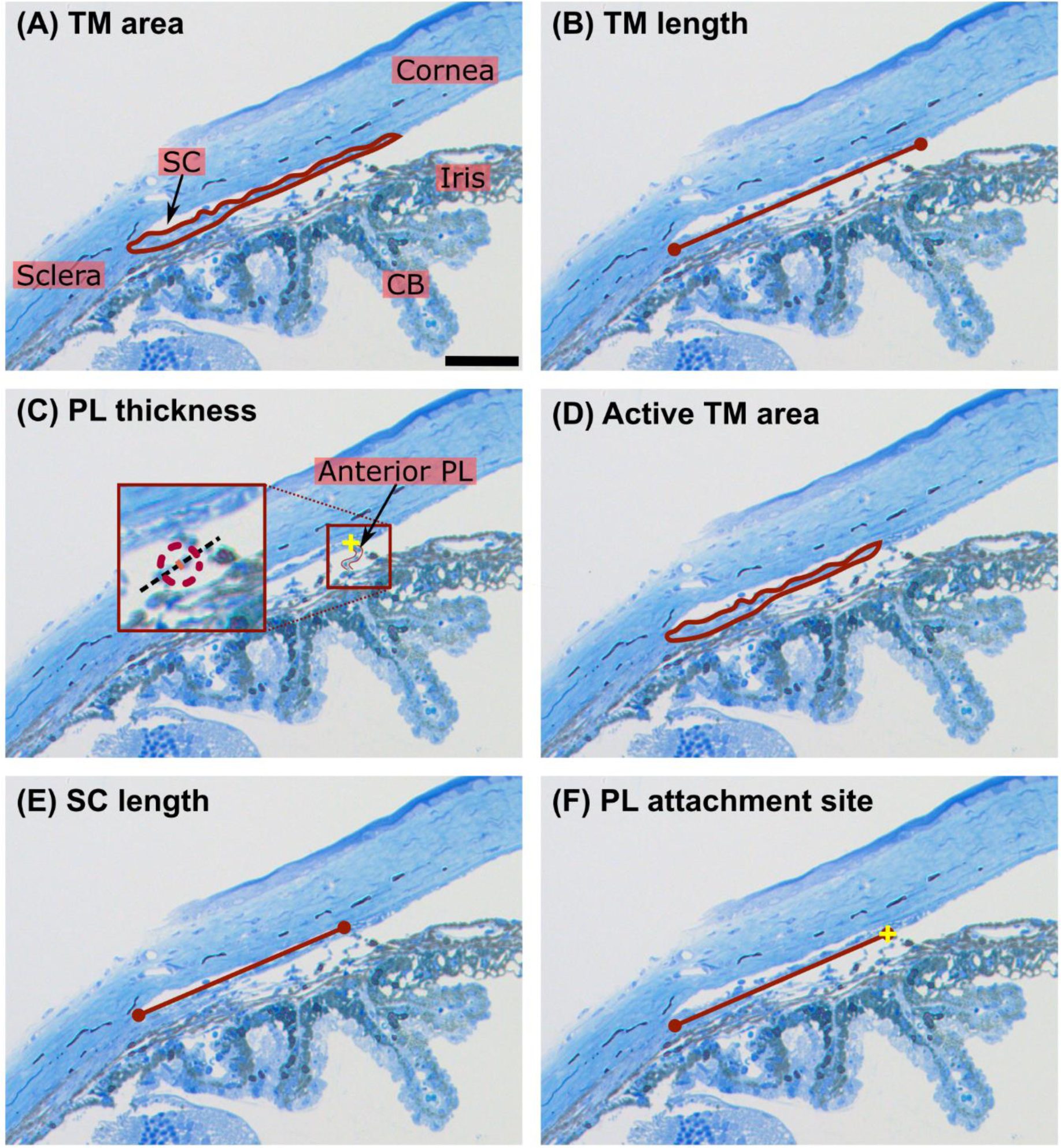
A representative sagittal section (from a young mouse) including the conventional outflow pathway tissues, showing definitions of the annotated features (red lines/boundaries) for morphometric characterization. Note the yellow cross in panel C, indicating the most anterior PL’s attachment site, which is the same location used to perform the measurement in panel F. Additionally, in panel C, the identified anterior-most PL is delineated, and in the expanded window, a black, dashed line is drawn along the local axis of the PL, i.e. in a direction perpendicular to the thickness measurement. TM = trabecular meshwork, PL = pectinate ligament, SC = Schlemm’s canal, CB = ciliary body. Scale bar = 50 µm.

The pectinate ligaments (PLs), as their name implies, form a comb-like fibrous structure located at the angle of the anterior chamber, bridging between the iris and the TM. PL structure varies substantially between species [20], suggesting potential evolutionary significance for this tissue. For example, piscine PLs (or annular ligaments) tend to be more extensive than in mammals [21,22]. Notably, in primates, including humans, the PLs are less extensive, with a porous presentation that is visible by gonioscopy.

Due to the less complex structure of the TM in mice, we hypothesize that the PLs play a role in mechanically stabilizing the TM and SC by imposing mechanical tension that resists the normal pressure differential from the anterior chamber to SC lumen. Our objective in this study was to test this hypothesis via a combination of histology-based morphometry (histomorphometry), *in vivo* optical coherence tomography (OCT) imaging, and computational modeling of the conventional outflow pathway tissues. All work was carried out on mice [23], a common model to study aqueous humor dynamics.

## Methods

We first histomorphometrically characterized key features of the primary outflow pathway in mice of different ages. We then augmented this histomorphometric data with OCT imaging to create age-specific computational (finite element [FE]) models of the murine outflow pathway from mice of different ages. Finally, we conducted forward FE analyses to investigate the effect of PL structure and mechanics on the biomechanical response of TM/SC to IOP as a function of age. The experimental procedures are described in detail elsewhere with a specific focus on morphological and biological effect of aging on aqueous outflow parameters [23]. Here, we pursue an in-depth analysis of morphology and biomechanics of the PLs.

### Animals

All the experimental procedures were approved by the Institutional Animal Care and Use Committees (IACUC) at Duke University and followed the tenets of the Association for Research in Vision and Ophthalmology (ARVO) Statement for the Use of Animals in Ophthalmic and Vision Research. Three groups of wild-type C57BL/6 mice were used in this study, as reported elsewhere [23].

- Group 1 was used for histomorphometry and consisted of eight young (2-3 months), six middle-aged (12-13 months), and seven elderly (29 months) mice. All mice for histomorphometry were male except for two females in the young cohort.
- Group 2 was used for high spatial resolution OCT imaging of the TM and SC (“TM-OCT”) and consisted of five young (4-6 months), seven middle-aged (11-16 months), and five elderly (25-31 months) mice. These mice were all male except for three females in the young cohort and two mice of unknown sex in the middle-aged cohort.
- Group 3 was used to evaluate IOP-induced iridial deformation via OCT imaging of the anterior segment (“Iris-OCT”). The young (3-6 months), middle-aged (9-13 months), and elderly (29-32 months) cohorts included five, seven, and five mice, respectively. In the middle-aged and elderly cohorts, all animals were male except for two females in the middle-aged cohort, while in the young cohort, all animals were female except for one male.

Please note that the age grouping used in this study essentially corresponded to those reported in our related paper [23].

### Histomorphometric analysis

Not all relevant anatomic features were visible on OCT images. Thus, we used sagittal histological sections through the outflow tract of mice in Group 1 to obtain some of the anatomical information needed to create the FE models. Anesthetized mice were euthanized by decapitation, and the eyes were immediately enucleated and immersion-fixed in 4% paraformaldehyde + 1% glutaraldehyde at 4 °C overnight. The eyes were then bisected, and the posterior segments and lenses were discarded. The anterior segments were cut into four quadrants and each quadrant was embedded in Epon, after which 0.5 µm thick sagittal sections were cut using a microtome, stained with methylene blue, and imaged (Axioplan2, Carl Zeiss MicroImaging, Thornwood, NY, USA; image resolution of ca. 0.26 μm/pixel) to visualize the outflow pathway tissues (Figure 1). For further details of these experimental procedures, please see Li *et al.* [23–25].

We manually identified and segmented the following features of interest from the resulting images (Figure 1):

- TM cross-sectional area (“TM area”), defined as the area of TM tissue bounded by the posterior terminus of Descemet’s membrane and by the most posterior point of SC (Figure 1A).
- TM length, measured as the distance between the two (anterior-posterior) extremes of the TM, measured using a straight line (Figure 1B).
- PL thickness, obtained by first identifying the most anterior PL that visibly connects the TM and ciliary body/iris, even if partially discontinous, and then measuring its thickness perpendicular to the local axis of the PL (Figure 1C).
- Active TM cross-sectional area (“Active TM area”), defined as the area of TM tissue lying directly adjacent to the lumen of SC (Figure 1D). Note that “Active TM area” is ≤ “TM area”.
- SC length, measured from the most posterior to the most anterior point of the SC lumen using a straight line (Figure 1E).
- PL attachment site(s), measured as the straight-line distance parallel to the line defining the SC length from the posterior end of SC to the identified PL from Figure 1C (Figure 1F).

Given the subjectivity in identifying some of these features, the segmentations were independently carried out by four separate annotators who were masked to mouse age and sample identity. All segmentations were performed using ImageJ (Fiji distribution; various 2023 versions [26]). The results were then compiled, and the annotators met to discuss the segmentation results and resolve any disagreements. The annotators then performed a second round of segmentation under the same masking conditions. After the second round of segmentation and discussion, three images were removed from the data set (two from the young cohort and one from the elderly cohort) because the tissues could not be reliably segmented. As a result, a total of 6 histological images were analyzed per age cohort, corresponding to 24 possible measurements (4 annotators × 6 images) per feature of interest. We then calculated the average “active TM thickness” in each image by dividing, for each annotator, the active TM area by the corresponding SC length, generating an additional 24 values per age cohort. Of note, due to the complex structure of the outflow tissues and inter-annotator variability, there were several unsuccessful annotations in some images. For instance, in the young cohort, a PL was not discernible in several samples (13 unmeasurable PL widths out of a total of 24), suggesting under-development of PLs in young animals (see the *Discussion* section).

As mentioned above, it was necessary to combine the morphometric data with both OCT and histologic images to generate the FE models. We reasoned that such a combination would be most robust if we scaled all dimensions by a common length apparent in both OCT and histologic images. We selected SC anterior-posterior length (“SC length”) as the common feature for this purpose. Hence, we calculated two ratios from histologic images: the “TM ratio,” defined as (TM length)/(SC length), and the “PL ratio,” defined as (PL attachment distance)/(SC length). We accounted for age variation by using age-specific TM ratios, i.e., we computed the ratio of average TM length divided by the average SC length for each age cohort, and we repeated the same process for the PL ratio. These ratios were later used in creating the FE models (see *Model geometry creation and FE mesh generation* section, below).

### OCT Imaging

Mice were anesthetized using an intraperitoneal injection of ketamine (100 mg/kg) and xylazine (10 mg/kg). A drop of 0.5% proparacaine was applied to the eye [23]. Animals were secured on a custom-made platform, and a single pulled glass microneedle filled with phosphate-buffered saline was inserted into the anterior chamber of one eye. The microneedle was connected to both a manometric column to adjust IOP and a pressure transducer to continuously monitor IOP levels using LabChart software (ADInstruments, Colorado Springs CO).

Two types of OCT imaging were performed using a high-resolution spectral domain OCT system (SD-OCT Envisu R2200, Bioptigen Inc., Research Triangle Park, NC, USA). In the first type of OCT image (TM-OCT, Group 2), we acquired sagittal scans focused on the iridocorneal angle, including the TM and SC lumen, at IOPs of 10, 15, and 20 mmHg. We identified the iridocorneal angle structures following our previously established techniques [25]. In the second type of OCT image (Iris-OCT, Group 3), the OCT scan head was placed near the corneal apex to acquire images of half of the anterior chamber at IOPs of 10, 15, and 20 mmHg. Additional IOP steps were also used but were not needed in this study [23,27].

### Model geometry creation and FE mesh generation

We created age-specific models of the conventional outflow pathway tissues by combining the morphometric measurements and the TM-OCT images. Due to physiological relevance and minimal perfusion-induced deformation, the TM-OCT images from 10 mmHg served as the reference state for creating all models (Figure 2A, D, and G). For each age-specific model, we delineated the corneal and iridial margins in one representative TM-OCT image (Figure 2B, E, and H) using the 2D spline function in SolidWorks (version 2022 Dassault Systèmes, Waltham, MA, USA).

**Figure 2:**
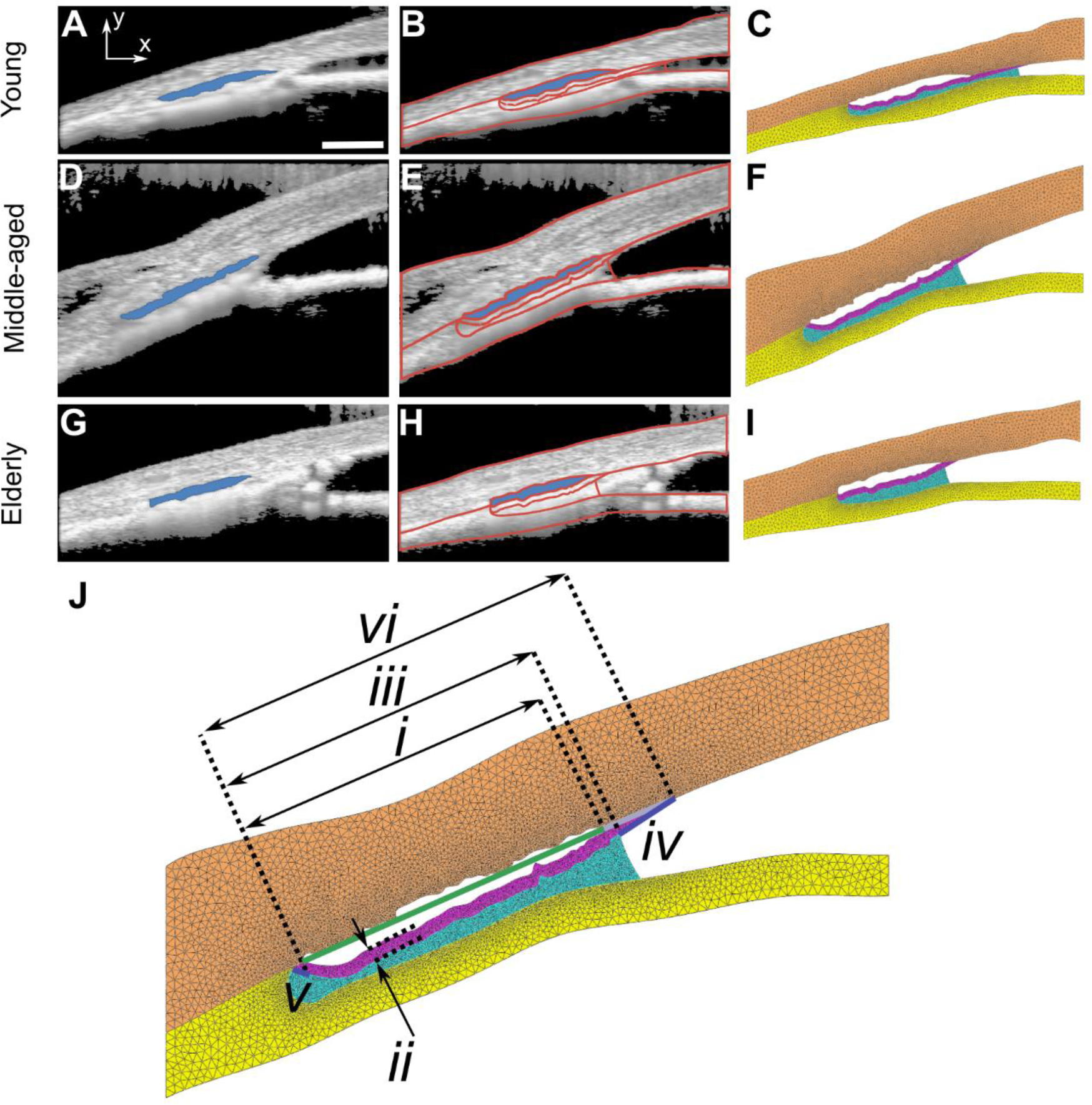
TM-OCT images, outlined features, and FE meshes for the (A-C) ‘young’, (B-F) ‘middle-aged’, and (G-I) ‘elderly’ models. Panels A, D, and G show OCT images acquired at an IOP of 10 mmHg with the lumen of SC overlain in blue. Panels B, E, and H show key structures segmented from OCT images or deduced from histomorphometric analysis (red overlaid curves), while panels C, F, and I show the resulting FE meshes. (J) Demonstration of a set of features used to create models, with emphasis on features based on histomorphometric analysis, including: (*i*) the line connecting the posterior and anterior ends of SC, (*ii*) an offset to create the inner margin of the TM based on the contour of the inner wall of SC, (*iii*) the agespecific calculated distance of the posterior end of SC to the anterior-most PL attachment site, (*iv*) a linear interpolation of the TM inner margin from the PL attachment site to cornea, (*v*) a linear interpolation of the inner TM margin connecting the posterior end of the TM to the apex of the irideocorneal angle, and (*vi*) the age-specific calculated TM length. Colors in C, F, I and J: orange = cornea and sclera; yellow = iris and other uveal tissue; blue = pectinate ligament region; purple = TM. Scale bar = 100 µm.

The details of the iridocorneal angle tissues were not resolved in these OCT images, and thus, as mentioned earlier, we relied on histomorphometry to reconstruct details of the outflow pathway tissues as follows. Using the segmented SC lumen, we created the TM by assuming that the TM’s inner margin was offset from the inner wall of SC by the age-specific average active TM thickness (Figure 2J, feature ‘*ii’*). This offset distance was measured normal to the line defining SC length (Figure 2J, feature ‘*i’*). Then, to complete the TM profile, we connected the ends of this inner TM margin to the cornea and root of the iris using linear extrapolations (Figure 2J, features ‘*iv’* and *‘v’*, respectively). In the young model the calculated PL and TM ratios were almost identical, and thus we extended the calculated TM length by c. 5% to facilitate meshing, which we note is well within the uncertainty in TM dimensions (Table 2). Finally, we used a constant TM thickness (averaged active TM thickness), rather than a spatially-variable thickness, as a practical approximation to keep the model tractable.

We modeled the PLs as a diffuse (distributed) material rather than attempting to model the complex, interconnected, and networked structure of the mouse PLs [14,28]. We recognize that this homogenized approach to modeling the PLs is an approximation, and we therefore created two additional models to assess its validity. In the first, we did not include the PLs at all, while in the second we only included a single PL “beam” (the anterior-most PL, as identified in Figure 1C), as previously described [25]. For consistency with previous work, we used the young model for these additional simulations (see supplementary Figure S2 for details). In both of these models, SC collapsed at low IOPs (IOP < 11 mmHg; Figure S2 D), which is contradictory to observations during OCT imaging [23,25]. This simple result suggested that treating the PL network as a diffuse (distributed) material, coupling the iris and TM, was the most physiologically-relevant option for our study, as discussed below.

The “distributed” PL material filled the iridocorneal angle, extending from the apex of the angle to the anterior-most PL attachment site (Figure 2J, feature ‘*iii’*), as determined for each age model using the “PL ratio” based on the histomorphometric analysis (Figure 1C and 2). The PL region was bounded by the interior surface of the TM and the anterior surface of the iris.

The final modeling step was to create a 3D CAD model by extruding the cornea, iris, TM, and PL as separate bodies, each 10 µm in height. This approach was preferred over creating an axisymmetric model due to the plane strain assumption (see the *Boundary conditions* section for further details).

To generate the FE mesh, we exported the CAD model as a STEP file (ASCII version 2) which was then imported into the Gmsh meshing software (version 4.11.1; [29]). We tessellated the volume using TET10 quadratic elements with overall mesh density set at 7.5 µm, confirmed to provide adequate numerical accuracy in a preliminary mesh refinement study. To enhance numerical accuracy, we used higher mesh densities in select locations. Specifically, we decreased the elemental characteristic edge length in the TM by a factor of 4 and in the PL region by a factor of 8. We used the average first principal Lagrangian strain (Ξ_I_) and SCLA estimates for the most compliant case from each age model and ensured that the error relative to an exhaustively refined mesh was less than 1% for Ξ_I_and SCLA (relative to the baseline SCLA of the corresponding age model). The meshes were generated using the Frontal algorithm, and ten steps of smoothing and optimization were conducted to enhance mesh quality [30].

### Boundary conditions

We used a plane strain modeling approach, justified by the conventional outflow region’s small size relative to its distance from the anterior chamber’s axis of symmetry. Further, we constrained the posterior (scleral) end of the model and the anterior corneoscleral surface (Figure 3), which was justified based on the significantly higher stiffness of the sclera and cornea relative to the key tissues of our model (see *Tissue constitutive models* section below). In addition, the right (central) edge of the cornea and iris were constrained to move only in the ‘y’ direction (Figure 3), which allowed for anterior chamber deepening during IOP elevation. Increasing IOP caused posterior displacement of the iris, as observed from the OCT images of the anterior chamber (see supplementary Figure S1). We used the lower-resolution OCT images (Iris-OCT) to extract iris deformations, which we then applied as age-specific boundary conditions in the FE model. These OCT scans were from a different set of animals (Group 3) than the high magnification OCT images (TM-OCT; Group 2) due to technical difficulties in maintaining a consistent OCT field of view at all IOP levels and in aligning the field of view with the anterior chamber’s central axis in the high-magnification TM-OCT scans. Therefore, we measured the iris displacement along the ‘y’ axis of Iris-OCT images (Figure S1; same axes definition as in Figure 2A for TM-OCT images) at IOP levels of 15 and 20 mmHg relative to the configuration at 10 mmHg and prescribed this displacement in the FE model as an input (see supplementary Figure S1 for more details).

**Figure 3:**
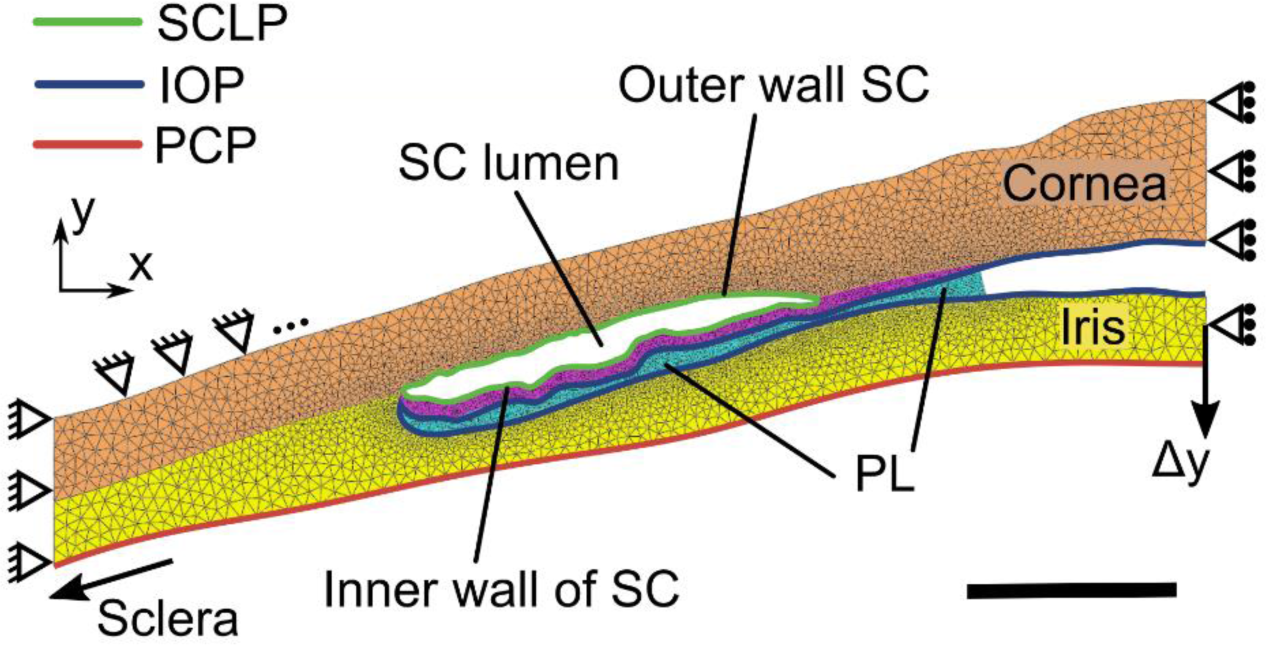
Boundary conditions for the FE model. Here, the young model is used as an example for illustrative purposes. Note the colored lines indicating the location where pressure loads were applied: SCLP = SC lumen pressure, IOP = intraocular pressure, PCP = posterior chamber pressure, and Δ*y* = prescribed displacement of the iris. Pinned support means no displacement in the ‘x’ and ‘y’ directions; rolling support indicates no displacement in the ‘x’ direction. Colors of individual sub-regions are the same as in Figure 2. Scale bar = 100 µm.

The pressure in the anterior chamber was specified based on the IOP levels imposed experimentally. For numerical modeling, IOP values were specified relative to the baseline pressure of 10 mmHg, i.e., we imposed a relative IOP of 5 mmHg (corresponding to 15 mmHg absolute) and of 10 mmHg (corresponding to 20 mmHg absolute). We set the posterior chamber pressure (PCP) to zero (10 mmHg absolute) based on the results of a prior study evaluating PCP as a data-fitting variable, which showed that PCP remained at 10 mmHg when IOP exceeded 10 mmHg [27]. Finally, SC lumen pressure (SCLP) was calculated based on IOP and the total resistance of the primary outflow pathway [25], which indicated SCLPs of 2 mmHg (12 mmHg absolute) and 4 mmHg (14 mmHg absolute) at IOP levels of 15 mmHg and 20 mmHg, respectively.

Finally, we imposed a contact boundary condition between the inner and outer walls of SC, which occurred when SC collapsed, using a node-on-facet algorithm and a penalty method to enforce contact. Specifically, we used an automatic penalty approach with the scaling factor set at 100 times TM stiffness.

### Tissue constitutive models

We used an incompressible neo-Hookean (“INH”) constitutive relation to model the cornea and iris.

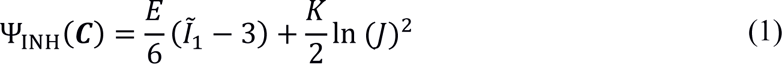

where Ψ_INH_is the Helmholtz free energy, ***C*** is the right Cauchy-Green strain tensor, 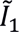 is the first invariant of the deviatoric right Cauchy-Green strain tensor, and *J* = det(***F***) is the Jacobian of the deformation gradient tensor. The stiffness *E* was set to 2700 kPa [25] for the cornea/sclera and 96.1 kPa for the iris [27]. Please note that although the term containing ln (*J*) is zero for a truly incompressible material, for numerical reasons it is necessary to include this term to enforce incompressibility in the finite element software. Towards this end, we set *K* (equivalent of bulk modulus) to 100 times the shear modulus to closely enforce incompressibility, as is standard [31].

Due to the fibrous structure of the PLs, we did not model them as incompressible; instead, they were modeled as a homogenized compressible neo-Hookean material (“NH”)

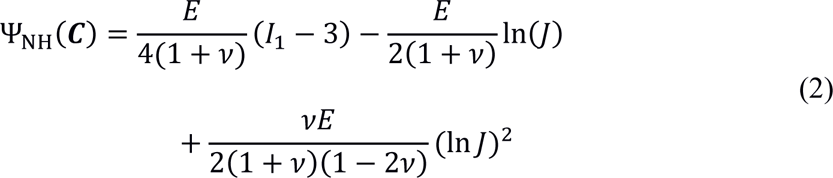

Here, *I*_1_is the first invariant of the right Cauchy-Green strain tensor, and *v* is Poisson’s ratio, which we set to zero due to the approximately parallel structure of the PLs noted above, assuming the PL beams behave as 1D elements lacking lateral contraction under axial tension. Therefore,

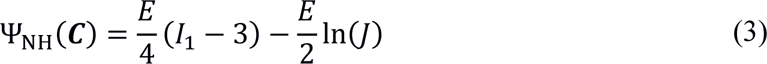

We assumed that the intrinsic stiffnesses of TM and PL were the same, based on their similar collagenous composition, but adjusted the stiffness value for the PLs to account for the high porosity of the PL network. In brief, we specified the effective stiffness of the PL relative to the TM using a linear relationship *E*_*PL*_ = β*E*_*TM*_, where β was the solid fraction of the PL network. We computed β from an analysis of scanning electron micrographs of rabbit PLs [20,32] since suitable high-resolution scanning electron micrographs from mouse eyes do not exist to the best of our knowledge, obtaining β = 2.3% (supplementary Figure S3). We then scaled this value down by a factor of 10 for mouse, since frontal sections through the mouse eye showed a more rarified PL network in mouse vs. rabbit [28]. The same value β = 0.23% was used for all ages.

### Forward finite element simulations and data analysis

We solved the FE models using FEBio (version 4.3.0, [33]), and developed a custom Python class to automate the simulations and post-processing. We specifically analyzed the deformations of aqueous outflow pathway tissues as a function of IOP in models with different TM and PL stiffnesses, for which a key metric was SC lumen cross-sectional area (SCLA). We varied *E*_*TM*_in a physiological range (5-150 kPa) [13], with *E*_*PL*_calculated from *E*_*TM*_as described above. We simulated 15 cases with varying *E*_*TM*_ and *E*_*PL*_.

To calculate SCLA, we used the *concave_hull* function (Shapely version 2.0.1, [34]) to fit a concave hull shape to the coordinates of the nodes on the inner and outer walls of SC (Figure 1). For consistency between models, we then computed a normalized SCLA, defined as

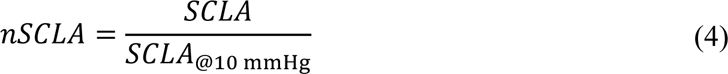

where *SCLA*_@10 mmHg_ is *SCLA* at the reference IOP (10 mmHg).

Finally, for each age-specific model and set of TM/PL stiffness values, we assessed SC lumen ‘patency’ using a quantity we term “Ultimate nSCLA,” defined as the value of nSCLA at IOP = 20 mmHg. An Ultimate SCLA value greater than one indicated a SC luminal area expansion at 20 mmHg relative to SCLA at 10 mmHg; conversely, Ultimate SCLA values less than one indicated an area reduction. In addition, as another measure of SC patency, we determined the lowest IOP at which SC ‘collapse’ occurred, which was defined as *nSCLA* < 33%. These definitions were based on informed estimations of the state of SC as it pertains to its capability to function within normal aqueous outflow, where collapsed regions of SC would not be able to function normally.

### Statistical analysis

We used a linear mixed effects (LME) model to study the effect of age on the morphometric characteristics of the mouse outflow pathway. Specifically, we used ‘age’ as the fixed effect and ‘annotator’ as a random effect. The significance threshold (α) was set at 5%. We also conducted a *post hoc* multiple comparison test on ‘age’ using Tukey’s HSD test with Bonferroni correction (critical α = 5%/3). In addition, we conducted a one-way ANOVA to test for the effect of age on TM- and PL-ratios (α = 5%), followed with *post hoc* Tukey’s HSD test (critical α = 5%/3). Statistical analysis was performed using JMP (Pro version 16, SAS Institute Inc., Cary, NC, USA)

## Results

### Morphometrical characterization of TM and PL in C57BL/6 mice

Histomorphometric analysis indicated that all measured dimensions changed with age, except PL thickness (Figure 4 and Table 1; *p* < 0.05 for the effect of age), with no effect of annotator (*p* > 0.05, Table 1). *Post hoc* multiple comparison tests indicated that, compared to the young cohort, the elderly cohort had a larger TM area (*p* = 0.014), Active TM area (*p* = 0.001), and SC length (*p* = 0.001) (Figure 4, Table 1, and Table S1). In addition, the middleaged cohort had a smaller SC length compared to the elderly cohort (*p* < 0.0001) (Figure 3, Table 1, and Table S1). Overall, there was a positive correlation between age and the size of the features.

**Figure 4:**
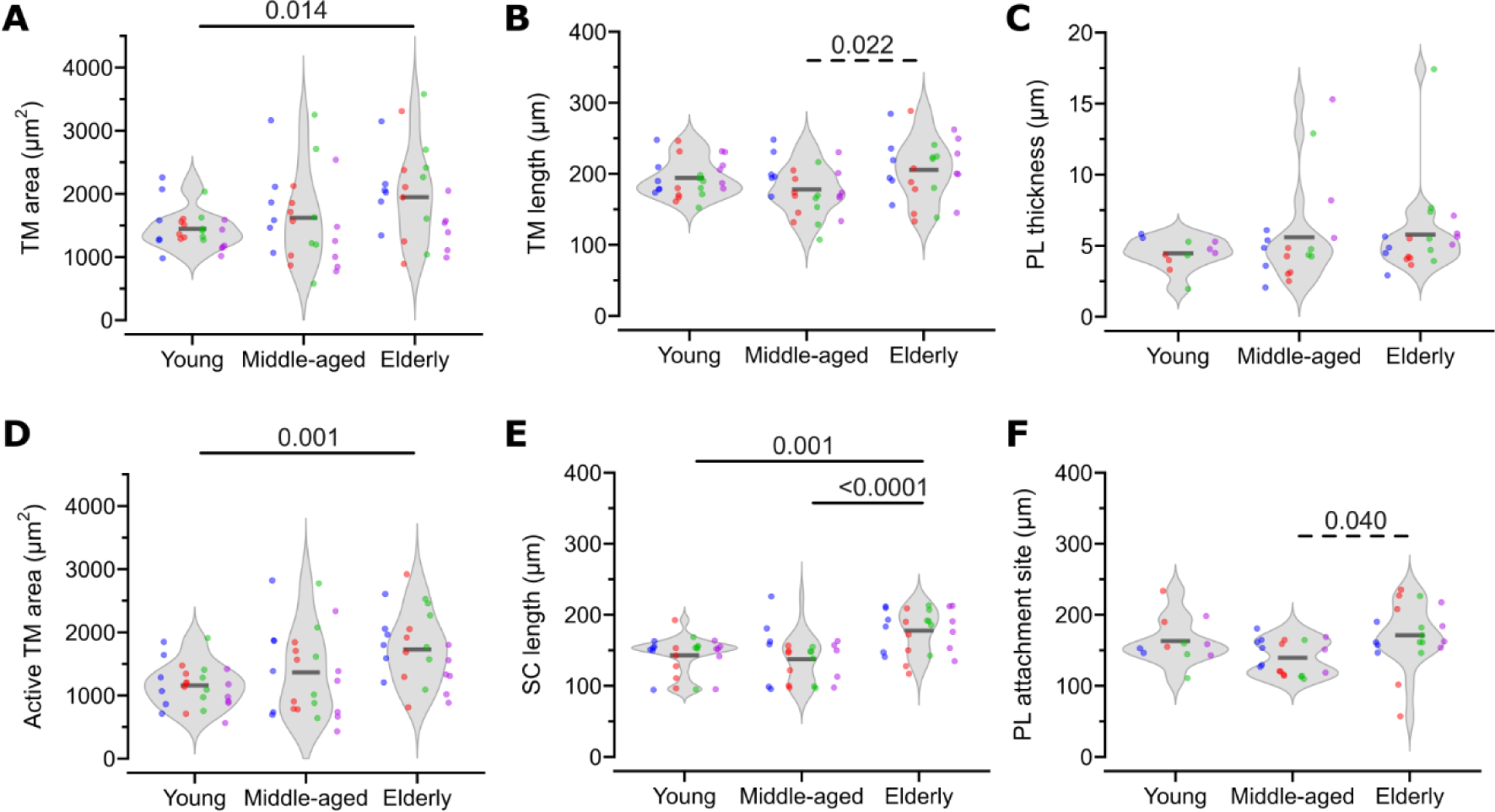
Results of the histomorphometric analysis of mouse outflow pathway tissue dimensions. For each feature, the data is shown using violin plots for each age cohort, with the mean indicated by a short black horizontal line. The results from different annotators are shown with colored dots, where each color corresponds to a unique annotator. Statistically significant differences by *post hoc* testing (after Bonferroni correction) are shown using solid horizontal lines (*p* < α/3 = 0.017), and trending differences are also shown using dashed horizontal lines (*p* < 0.05).

**Table 1:**
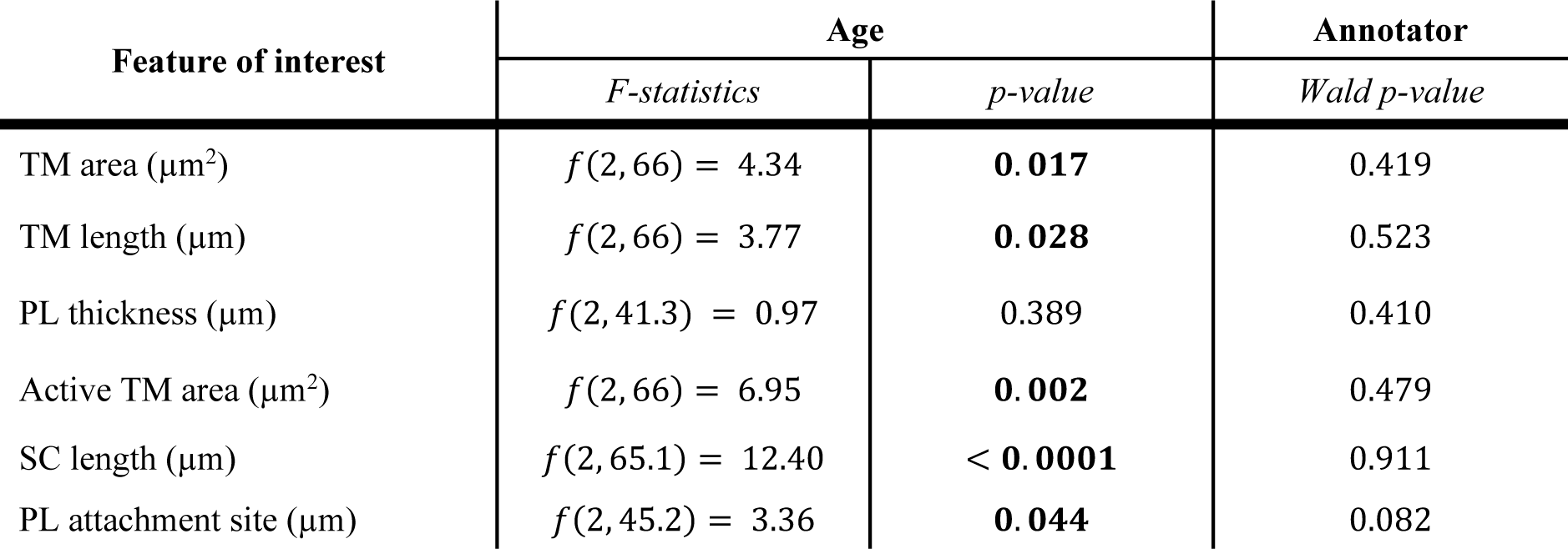
Results of the linear mixed effects (LME) model testing on the histomorphometric characterization of the C57BL/6 mouse outflow pathway. The columns provide F-statistics for the fixed effect (‘age’; **Bold**: *p* < 0.05), and Wald p-value for the random factor (‘annotator’), which tests the null hypothesis that the random effect was significantly different from zero (*Wald p-value < 0.05*) [56].

Furthermore, the TM- and PL-ratios were significantly affected by age (*p* < 0.05; Table 2). More specifically, the elderly mice had smaller TM- and PL-ratios relative to the young cohort (Table S3), indicating a moderately posterior localization of TM and PLs relative to SC lumen in elderly animals.

### Dependence of SC lumen area on TM/PL stiffness

All models (for all three age groups) demonstrated IOP-induced SC lumen collapse, as was also observed experimentally [23,25,35,36]. However, large TM/PL stiffness values caused unphysiological SC expansion at elevated IOP (Figure 5), which was not observed experimentally. More specifically, SC lumen area showed a highly nonlinear dependence on IOP and TM/PL stiffness in all age-specific models (Figure 5D). Interestingly, elevation of IOP did not guarantee reduction of SC luminal area. Instead, collapse only occurred at lower TM stiffnesses (Figure 5B, C, and D). At higher stiffnesses, SC lumen—counter-intuitively—expanded with increasing IOP (Figure 5A and D). Further, the dependence of normalized SC cross-sectional lumen area, nSCLA, on IOP was not always even monotonic. For instance, in the middle-aged model, there was an initial phase of nSCLA reduction with increasing IOP followed by a sudden expansion of the SC at *ca.* IOP = 11 mmHg (Figure 5D middle). But, as IOP further increased beyond 15 mmHg, SC luminal area gradually shrank. In a similar fashion, albeit more gradually, such behavior was observed in the elderly model, and to a lesser extent, in the young model (Figure 5D).

**Figure 5:**
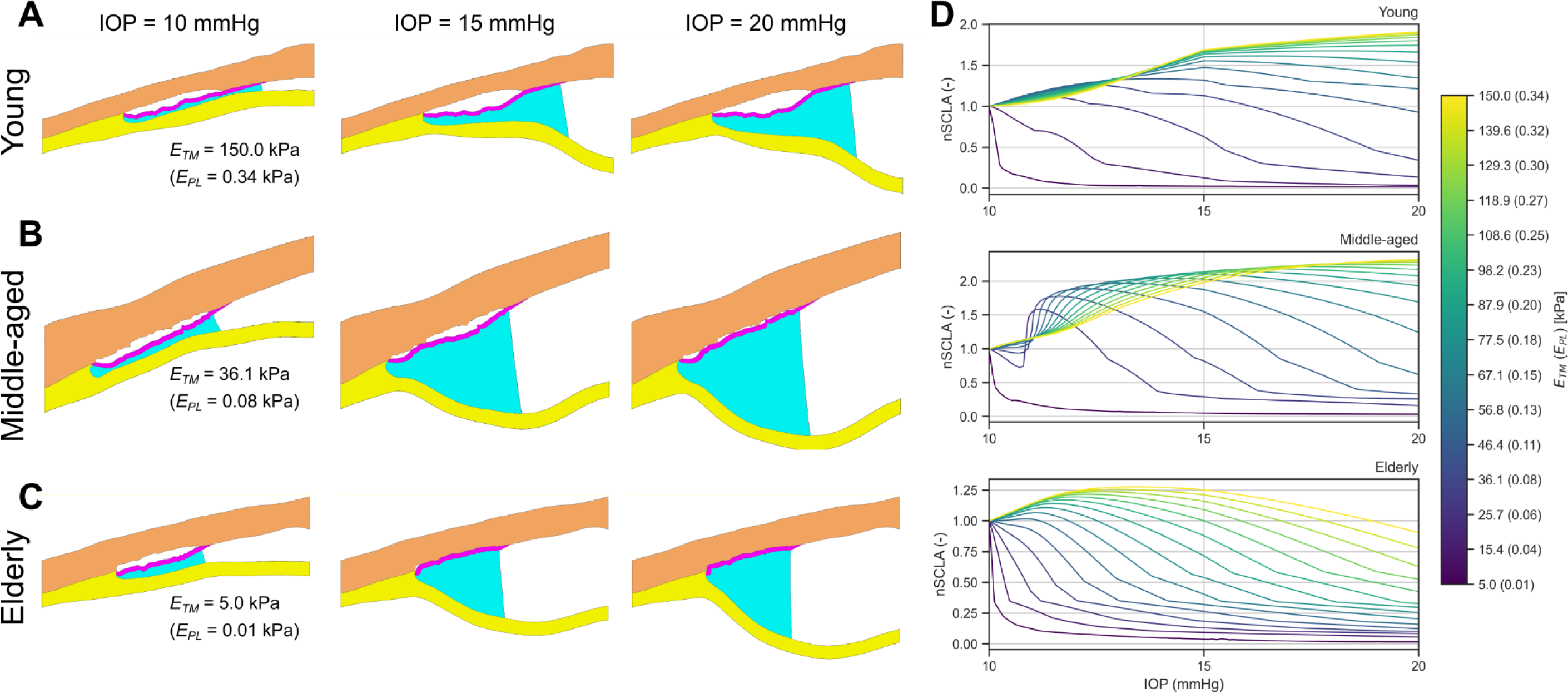
The results of forward FE simulations for each age-specific model, showing representative simulation results for the (A) young, (B) middle-aged, and (C) elderly models. In addition, (D) normalized SC lumen area (nSCLA; SC area normalized by the SCLA at 10 mmHg) vs. IOP is plotted for a range of E_TM values (color bar; 5 kPa ≤E_TM≤150 kPa). Interestingly, SCLA was not monotonically reduced by increasing IOP in the cases with higher E_TM, explainable by tension on the TM delivered by the PL due to the posterior deformations of the iris. Additionally, in the middle-aged simulations, there was an interesting rapid initial increase in SCLA followed by a gradual decrease, which was not evident in the simulations in the other age-specific models.

### Patency of SC lumen

To investigate the patency of SC, we evaluated: (i) the Ultimate nSCLA for each agespecific FE model vs. TM/PL stiffness (Figure 6A) and (ii) the IOP at which SC lumen first collapsed for each age-specific model vs. TM/PL stiffness (Figure 6B). Recall that in our approach, increasing TM stiffness also proportionally increased PL stiffness; thus, we simply denote stiffness as “TM/PL stiffness”. We observed that the Ultimate nSCLA strongly depended on TM/PL stiffness, with a lower stiffness giving a smaller Ultimate nSCLA (Figure 6A). In the young and middle-aged models, the Ultimate nSCLA was greater than 1 (expansion of SC lumen) for cases with *E*_*TM*_ > 46.4 kPa (*E*_*PL*_ > 0.11 kPa) in the young model, and *E*_*TM*_ > 67.1 kPa (*E*_*PL*_> 0.15 kPa) in the middle-aged model (Figure 6A). For the cases where the Ultimate nSCLA was smaller than 1, which are physiologically relevant, the elderly model showed a smaller Ultimate nSCLA for any given *E*_*TM*_compared to both young and middleaged models.

**Figure 6:**
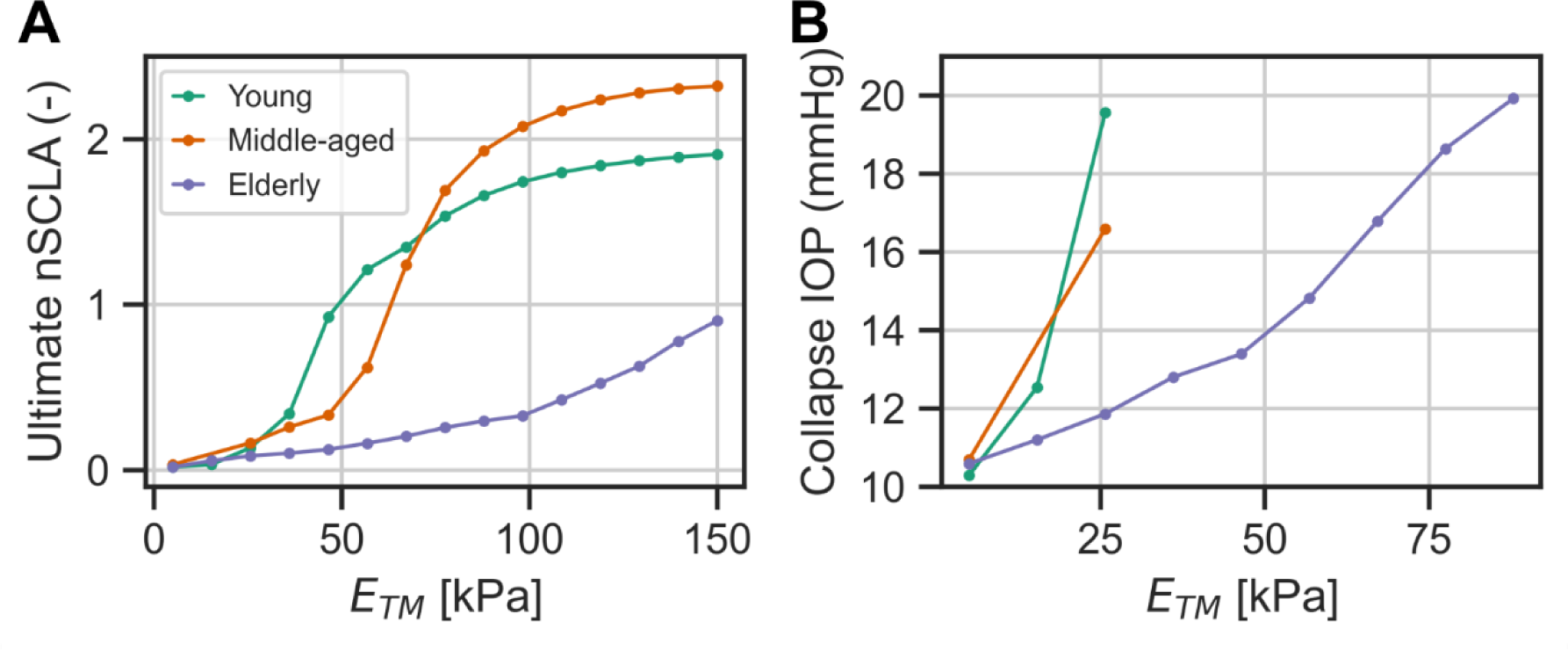
The patency of the SC lumen was characterized in each age-specific model by evaluating (A) the dependence of Ultimate nSCLA (nSCLA at 20 mmHg) on TM stiffness, and (B) the IOP at which SC lumen collapsed vs. TM stiffness. Note that in the young and middle-aged models SC collapse (B) did not occur in cases with stiffness higher that 26 kPa, which is the reason for the termination of the associated curves at that point.

Further, when investigating the collapse IOP for different TM/PL stiffnesses (Figure 6B), the highest *E*_*TM*_at which SC collapsed in the elderly model was *E*_*TM*_ = 87.9 kPa (*E*_*PL*_ = 0.20 kPa), which was higher than that of the young and middle-aged models *E*_*TM*_ = 25.7 kPa (*E*_*PL*_ = 0.06 kPa; Figure 6B). As expected, all models collapsed at a low IOP of 10∼11 mmHg at the lowest simulated *E*_*TM*_stiffness of 5 kPa (*E*_*PL*_ = 0.01 kPa; Figure 6B). Yet, surprisingly, for the same TM/PL stiffness levels, the elderly model collapsed at the lowest IOP compared to both other models (Figure 6B), except for the limiting case of *E*_*TM*_ = 5 kPa (*E*_*PL*_ = 0.01 kPa), where the models had a similar collapse IOPs.

## Discussion

In this study, we investigated the role of the pectinate ligaments (PLs) in stabilizing the conventional outflow pathway in the mouse eye, a ubiquitous model for studying aqueous humor dynamics. We treated PLs as a tissue with a biomechanical function and mechanical loading environment different from that of the TM. Unfortunately, because there are no direct measurements of PL biomechanical properties, we were forced to estimate their biomechanical properties from those of the TM. We showed that the stiffness and morphology of the PLs play a significant role in stabilizing the outflow pathway, and hence, in supporting the TM, ensuring patency of SC lumen, and in influencing how outflow pathway tissues deform as IOP changes. These observations reveal a previously unexplored role for the PLs in murine aqueous humor dynamics. Indeed, in our computational models in which the PLs were not present, SC was predicted to collapse at very low IOPs, in opposition to what is observed experimentally. Considering the anatomy of the mouse TM (few beams and proportionately much thinner in the interior-exterior direction than in larger eyes), it is perhaps not surprising that mechanical stabilization of the TM in the mouse is needed to avoid complete SC collapse, which would cause partial loss of functionality of the primary aqueous humor outflow pathway and significant elevation of IOP.

It is interesting that in one of the few studies of PL mechanics [37] reported that severing the PLs in a dog led to an expansion of the TM and a corresponding increase in outflow facility, i.e. opposite to our predictions. This contradiction is likely due to a species effect: dogs have a more robust TM and iris that may be more resistant to collapse and bowing, respectively, so that the canine PLs likely play less of a TM stabilizing role than in the mouse, but rather serve to maintain the angle between the TM and iris. Consistent with this interpretation, canine PLs appear to have a subtle, but important difference in morphology vs. the mouse: in the dog, the PLs attach to the iris immediately proximal to the root and then run anteriorly so that their anterior-most attachment point lies in the corneal limbal region rather than in the TM (see Figure 2 from Morrison and Van Buskirk [37]). The net effect is that the course of the PLs in the canine eye is more parallel to the beams of the TM than in the murine eye, where there seems to be a more interior-exterior alignment of the PLs leading to iris bowing directly pulling the TM interiorly. We conclude that inter-species differences in PL morphology and function may be important and merit future attention.

One interesting observation was the non-monotonic dependence of Schlemm’s canal luminal area on IOP (Figure 5D), which illustrates the complex biomechanics of the outflow pathway. This behavior was particularly evident in the middle-aged model (Figure 5D, middle) which illustrates two opposing factors impacting the configuration of outflow tissues: (i) IOP elevation tends to collapse SC lumen by outwards displacement of the TM, whereas (ii) posterior iridial movement with increasing IOP tends to pull the TM inward and open SC. The complex shapes of the specific curves in Figure 5D depend on TM stiffness, PL stiffness, and geometric factors such as TM thickness and PL anterior-posterior extent (Figure 5 A-C). The complexity of these effects is reinforced by observing the slope discontinuities in the nSCLA vs. IOP curves, which were due to progressively larger contact between the inner and outer walls of SC. In future work, we should also investigate the role of septae in the SC lumen [38], known to be present in human eyes but largely uninvestigated in the murine eye.

Although PL biomechanics has not been the focus of many investigations, PL morphology has been studied in a variety of domestic mammals, including dogs, horses, and rabbits, via scanning electron microscopy (SEM) and other types of histology [20,21]. PL morphology varies between species via differences in PL width, length, spacing, and insertion angle into the TM. Furthermore, PL morphology also varies locally within the same eye. One factor suggested to influence PL structure is age [20]. While the effect of aging on PLs was not a primary focus of this work (see [23]), our models were based on images from different aged animals. Therefore, it is worth exploring our results from that perspective.

Our histomorphometric analysis demonstrated a dependence of structural features on age (Figure 2 and Table 1) with certain features becoming more prominent with age (Figure 2), as expected. For instance, in the young cohort, the PL was relatively underdeveloped. In fact, we were unable to observe PLs in 13 out of 24 annotations from young eyes. Once developed, however, our results only showed a modest, although not statistically significant, increase in PL thickness with age. Due to the small thickness of the PLs, there were challenges in accurately measuring their thickness from light micrographs. More accurate measurements of the PL would enable us to assess evaluate PL thickness with age. Further, the PLs are 3D structures, and our 2D analysis is likely inadequate to fully evaluate the microscopic structure of the PLs. Nonetheless, we hypothesize that there is an association between age and PL structures that warrants future investigation. In addition, our analysis showed that age affected the TM and PL ratios (Table 2), with older eyes having larger SCs, and thus, smaller TM and PL ratios (Figure 2).. We hypothesize these differences might be related to development and growth of more prominent PLs with age, thereby providing more support to the TM, preventing SC collapse in middle-aged and older eyes.

**Table 2:** Ratios of the features measured from histomorphometric analysis used for creating the FE models. (**Bold**: **\****p* < 0.05)

|  | *Young* | *Middle-aged* | *Elderly* | *F-statistics* | *p-value* |
| --- | --- | --- | --- | --- | --- |
| TM-ratio<br>(TM length/SC length) | 1.40<br>± 0.26 | 1.35 ± 0.43 | 1.16 ± 0.19 | $f(2, 68) = 3.86$ | **0.026\*** |
| PL-ratio<br>(PL attachment site /SC<br>length) | 1.36<br>± 0.46 | 1.11 ± 0.36 | 1.00 ± 0.25 | $f(2, 45) = 3.83$ | **0.029\*** |

In addition, our simulations indicated a connection between iris deformation and TM/PL stiffness (Figure 5A-C), which was demonstrated by differences in the slope of the iris midline at the pupillary (right) edge of our models (IOP = 15 mmHg and 20 mmHg columns; Figure 5A-C). This observation suggests a connection between iris deformations and aging, which can influence the association between anterior chamber deepening (AC deepening) and IOP. We know AC deepening significantly affects outflow facility and can also artificially elevate facility in experiments [39–41]. Therefore, in perfusion preparations where the iris is intact, the association between AC deepening and changes in outflow facility may be explained by PL tension, where with more AC deepening the PLs open TM and prevent SC collapse (as observed in some cases in our study; Figure 5A), therefore increasing facility. Thus, it would be worth more closely exploring whether there are connections between TM/PL stiffness and iris deformations under, for example, various physiologically occurring iris contractions (e.g., at different light levels [42]), and at different ages to study connections between iris deformation, age, and outflow pathway function.

As a further age-dependent observation, our simulations indicated a strong effect of age on the biomechanical properties of outflow tissues. For example, by comparing modeled nSCLA vs. experimentally observed nSCLA at elevated IOP levels (IOP>10 mmHg), a significant stiffening of TM was observed with age [23]. Of note, our predicted TM stiffness for young mice (∼29 kPa [23]) was consistent with previous results on young mice [25]. We also predicted that elderly eyes had the smallest Ultimate nSCLA for a given TM/PL stiffness (Figure 6A). Overall, these observations indicate an increased resistance to IOP-induced deformations of outflow tissues with age, due to a combination of changes in biomechanical properties and tissue geometry. It is also important to note that the stiffening of TM/PL with age reduces the chances of SC collapse, yet, TM stiffening has been associated with glaucoma pathogenesis [12,43–46], which appears to be paradoxical. However, we would like to highlight that decreased susceptibility to SC collapse is different from changes in outflow facility due to TM stiffening. For example, suppose that the TM and PL both stiffen. Our model suggests that this would keep SC patent, and thus the flow resistance of the TM/SC would be dominated by the TM, since the flow resistance within the lumen of SC is generally much less than the resistance across the TM [47]. In this case, a stiffened TM would be associated with greater flow resistance, and so no paradox exists. Harder to understand is the case where both the TM and PLs are soft: here our model predicts that SC collapse could occur and thus lead to a large total outflow resistance even with a soft TM, which is indeed paradoxical. One possible explanation is that SC collapse is thought to contribute to ocular hypertension only at extreme collapse levels [48], and this effect is highly nonlinear in SC lumen size. Thus, so long as the PLs can partially prevent collapse, there may be no meaningful elevation of SC luminal flow resistance, so that total outflow resistance remains relatively low. We do not claim to entirely understand this, and we suggest that unraveling this complexity will require direct experimental measurements of PL stiffness in normal and ocular hypertensive mice.

On a technical note on numerical modeling, it is important to mention that we here used quadratic finite elements (TET10) that have superior performance in modeling incompressible materials [49] than the linear elements that we previously used (TET4; [25]). This element type was essential for making comparisons between different FE models in this study and should be used as for future modeling work in studying outflow tissue biomechanics. This difference in element type likely explains differences between our current results and those seen in earlier studies from our lab, e.g. [25], where we previously predicted that a single PL could stabilize the TM.

It is worth noting this study had some limitations. First, due to a lack of information about PL biomechanical properties, we were forced to estimate PL stiffness based on information about PL porosity in a rabbit eye which required several assumptions. Fortunately, we kept the PL to TM stiffness ratio consistent between models, thus eliminating variability in this ratio as a source of difference between models. Nonetheless, more direct estimates/measurements of PL biomechanical properties, while challenging, should be undertaken in future. Second, while we used a consistent processing protocol for the histological images, the tissues dimensions could have been subjected to changes during preparation (e.g. shrinkage) that affected the FE models. In future investigations, the use of imaging technologies that allow in situ imaging of the intricate structure of outflow pathway (e.g., VIS-OCT [50]) may help overcome this limitation. Third, OCT imaging of mouse eye is subject to some technical limitations: *i*) because of the diminutive size of the mouse eye, the manual positioning of the OCT scan head, and IOP-induced changes in ocular geometry, there was inevitable shifting of the relative position of the scan probe and the eye between scans of the same eye at different IOPs, and *ii*) OCT imaging is not able to capture all details of the angle structures, leaving us no choice but to use histomorphometry to “inject” those structures into FEM models based on OCT images. This process, together with the anatomic variability between mice, inevitably led to some mismatches between histologically-determined dimensions and the individual OCT images (e.g. Figure 2H). These technical limitations added variability to our analysis, which was addressed by using lower magnification Iris-OCT images for certain parts of the modeling process, and by using age-specific average histomorphometric feature dimensions. In the future, hardware to accurately position the OCT scan head, and improved resolution (penetration depth) to better image angle structures might improve the repeatability of high-resolution OCT images. Fourth, we used 10 mmHg as the baseline IOP for our FE models, which is a limitation since this baseline state is not actually stress-free; however, since we used a nearly linear constitutive relation (i.e., neo-Hookean) and a normalized measure (i.e., nSCLA) as our main outcome parameter, the effect of this modeling choice on conclusions is expected to be minimal. Fifth, the current lack of information on PL biomechanics makes it difficult to verify whether the PL deformations predicted in our study (Figure 5) would be physiologically tolerated without violating the elastic material assumption of our model. It is possible that the PLs demonstrate inelastic behaviors (e.g., structural damage, plastic deformation, and local failure) [51,52]. Of note, for our models’ most compliant case (*E*_*TM*_ = 5.0 kPa, *E*_*PL*_ = 0.01 kPa), the maximum first principal engineering strains (averaged over the entire PL region) were c. 26 (Young), c. 10 (Middle-aged), and c. 6 (Elderly). However, we note that at low pressures, the PLs adopt a tortuous configuration (Figure 1), which may allow the PL tissue region to experience large strains without individual PLs experiencing inelastic behavior. These aspects of PL biomechanics merit further investigation. Sixth, we based our analysis on one strain of mice (i.e., C57BL/6); however, PL in other strains, such as FVB/N, may demonstrate somewhat different behavior. For example, there is a different correlation between AC deepening/outflow facility vs. pressure in C57BL/6 as compared to FVB/N mice [53]. Hence, future investigations should consider other mouse strains (and species) to better elucidate the role of PL in aqueous outflow. Finally, although sex differences are likely important in glaucoma [27,54], due to the scarcity of female samples in our data set, we did not consider sex as a biological variable in our analysis. Therefore, caution is advised interpreting these results for both sexes.

In conclusion, our simulations showed that PL biomechanics plays an essential role in maintaining stability of the outflow pathway tissues in the mouse eye. In the future, it will be important to directly characterize PLs’ biomechanical properties, preferably by direct biomechanical testing of PL beams, e.g., using atomic force microscopy. We note that PLs are less prominent in healthy human eyes and may thus be less clinically important relative to the preclinical studies in mouse eyes. However, structures like pectinate ligaments can persist (see e.g. Figure 11 in [55]) in human eyes with developmental pathologies leading to congenital (infantile) glaucoma, and it would be of interest to evaluate their role in such cases.

## Conflict of interest

The authors declare no conflicts of interest.

## Acknowledgments

We gratefully acknowledge Prof. Darryl Overby of Imperial College London for his insightful comments regarding interpreting of our results. Further, we note the generous financial support from The BrightFocus Foundation (postdoctoral fellowship G2021005F, BNS), NIH (K99EY035360 [BNS], T32EY007092 [NSFG], R01EY030871 [AJF], P30EY006360, P30EY005722, R01EY031710 [CRE and WDS], and R01EY030124 [WDS]), the Alfred P. Sloan Foundation G-2019-11435 (NSFG), the Georgia Research Alliance (CRE), and Department of Veterans Affairs Rehab R&D Service Career Development Awards (AJF; CDA-2; RX002342).

## Supplementary Material

**Table S1:** Results of the multiple comparison Tukey’s HSD test on the ‘age’ factor on histomorphometric characterization of the C57BL/6 mouse outflow pathway. (*\*p<0.05, **p<0.017*)

| Feature of interest | Age comparison |  | *p-value* |
| --- | --- | --- | --- |
| TM area ($\mu\text{m}^2$) | Young | Middle-aged | 0.575 |
|  | Young | Elderly | **0.014\*\*** |
|  | Middle-aged | Elderly | 0.149 |
| TM length ($\mu\text{m}$) | Young | Middle-aged | 0.243 |
|  | Young | Elderly | 0.513 |
|  | Middle-aged | Elderly | **0.022 \*** |
| PL thickness ($\mu\text{m}$) | Young | Middle-aged | 0.445 |
|  | Young | Elderly | 0.417 |
|  | Middle-aged | Elderly | 1.000 |
| Active TM area ($\mu\text{m}^2$) | Young | Middle-aged | 0.386 |
|  | Young | Elderly | **0.001\*\*** |
|  | Middle-aged | Elderly | 0.055 |
| SC length ($\mu\text{m}$) | Young | Middle-aged | 0.809 |
|  | Young | Elderly | **0.001\*\*** |
|  | Middle-aged | Elderly | **< 0.001\*\*** |
| PL attachment site ($\mu\text{m}$) | Young | Middle-aged | 0.229 |
|  | Young | Elderly | 0.861 |
|  | Middle-aged | Elderly | **0.040\*** |

**Table S2:**
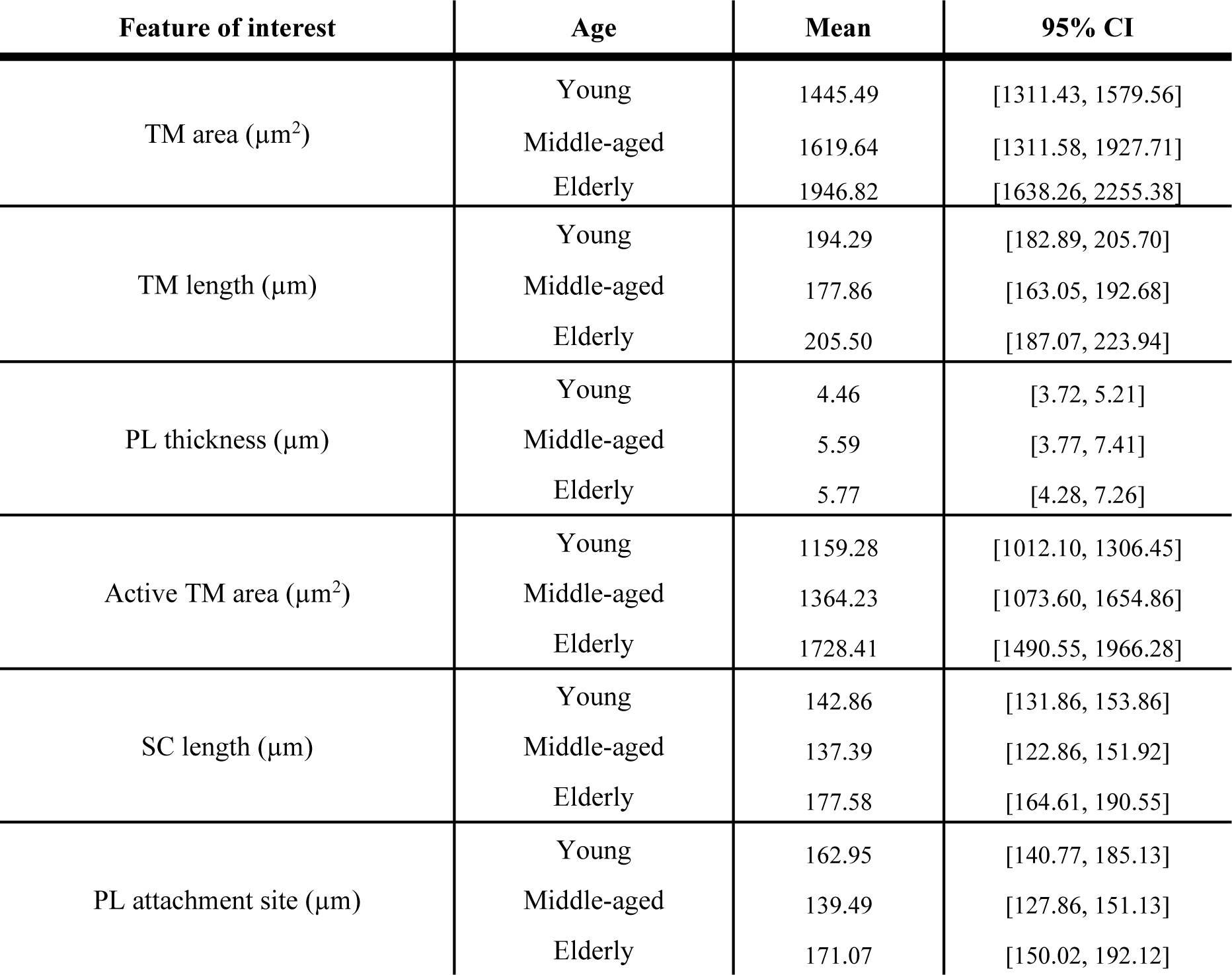
Summary statistics for the histomorphometric characterization of C57BL/6 mouse outflow pathway features by age cohort. Data is shown as mean and 95% confidence interval (95% CI).

**Table S3:** Results of multiple comparison Tukey’s HSD test on TM- and PL-ratios. (*\*p<0.05, **p<0.017*)

| Feature of interest | Age |  | *p-value* |
| --- | --- | --- | --- |
| TM-ratio | Young | Elderly | **0.030\*** |
|  | Young | Middle-aged | 0.094 |
|  | Middle-aged | Elderly | 0.887 |
| PL-ratio | Young | Elderly | **0.023\*** |
|  | Young | Middle-aged | 0.144 |
|  | Middle-aged | Elderly | 0.620 |

**Figure S1:**
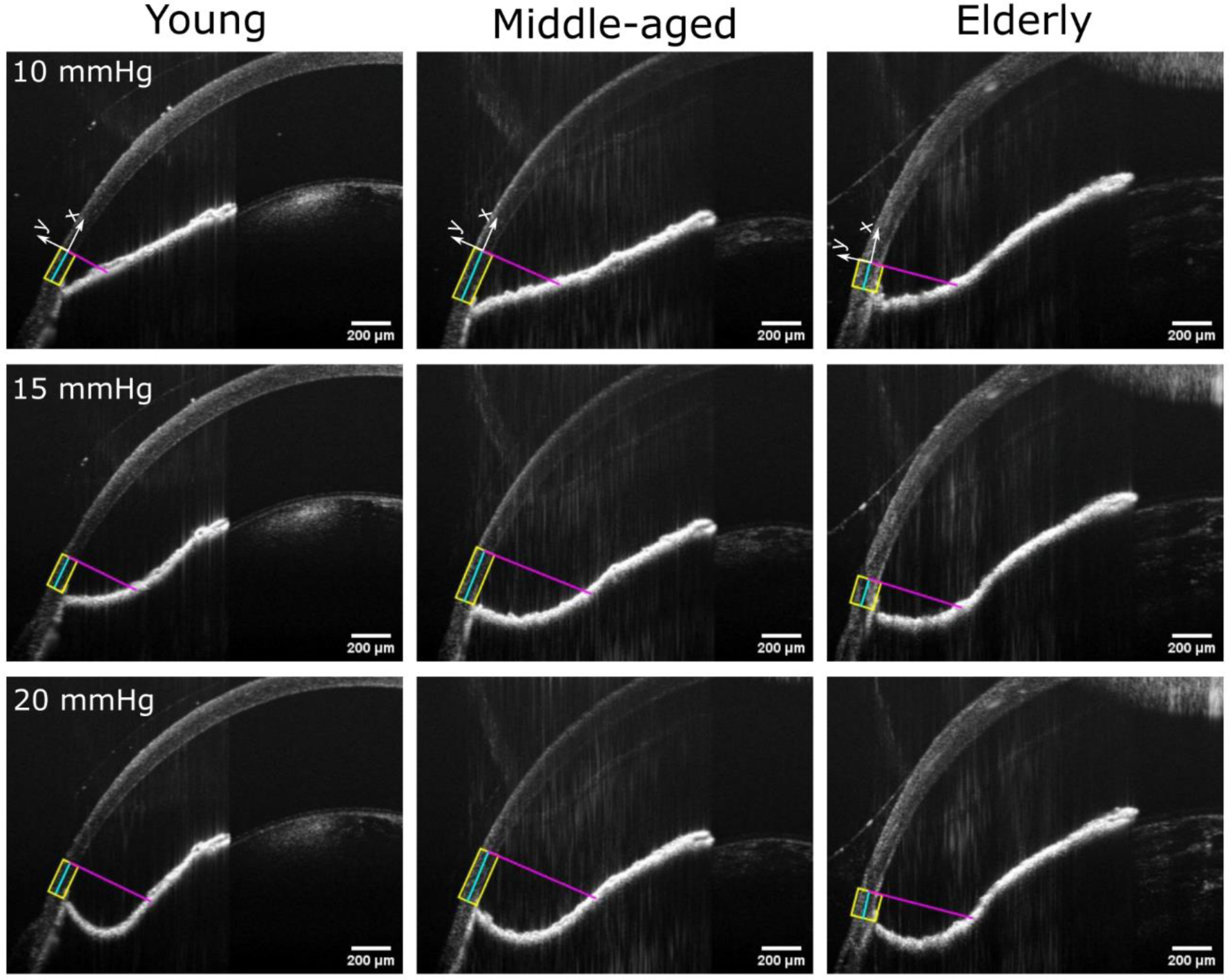
Iris-OCT images, used to evaluate the IOP-induced deformations of the iris. We drew a line segment (cyan) along the midline of the cornea (yellow lines), starting at the apex of the iridocorneal angle and extending to the edge of the cornea typically visible in TM-OCT images. We then constructed a perpendicular line segment connecting the end of the cyan line to the midline of the iris (purple). We determined the length of this purple line segment along the ‘y’ axis for different IOPs, and we evaluated iris displacement in the FE model based on the length change of this line segment. See top-row panels for the definition of the ‘y’ axis.

**Figure S2:**
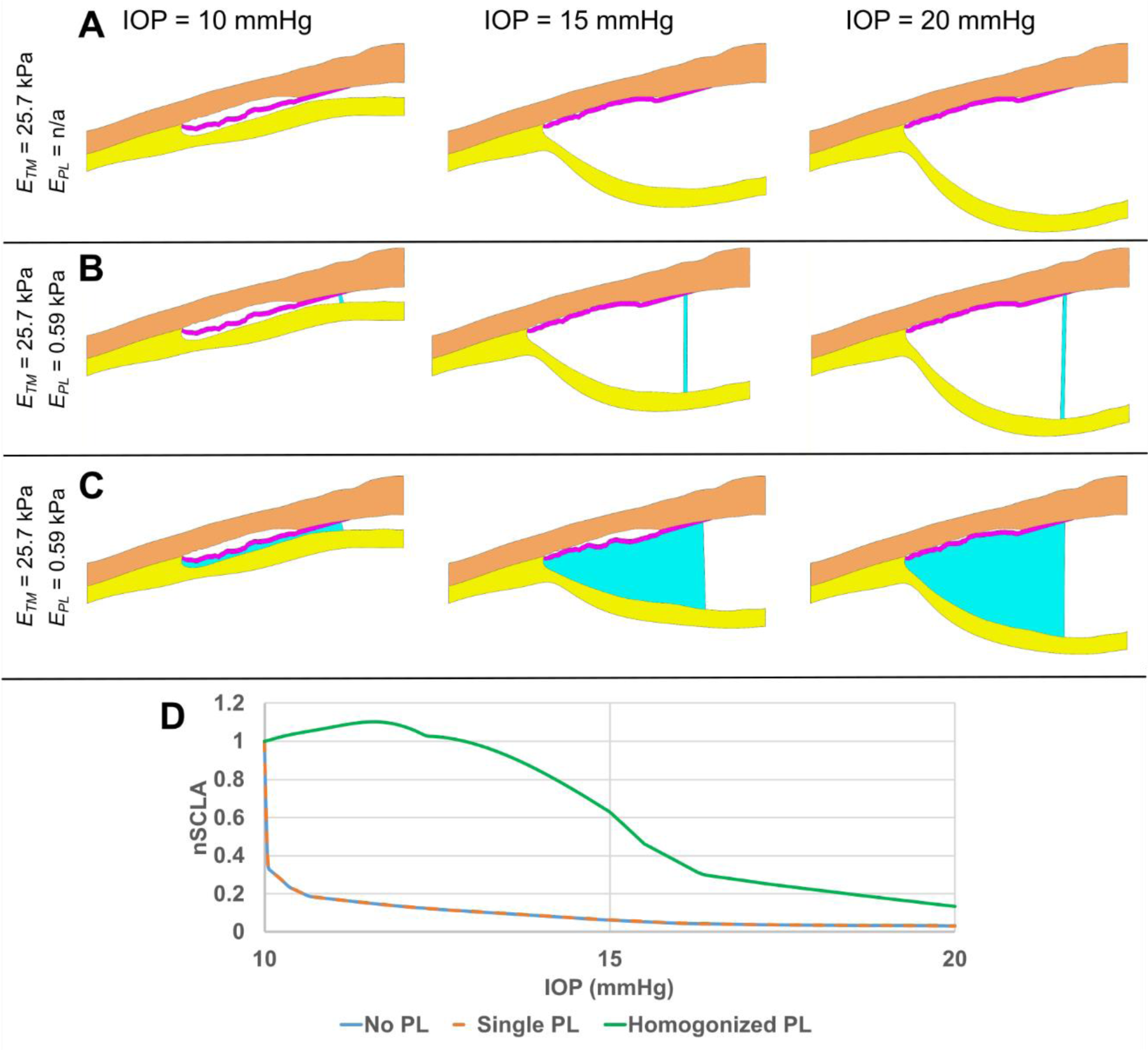
Computed tissue deformations vs. IOP in three cases with different PL structures. **(A)** No PL, **(B)** a single PL at the anterior end of the PL region, and **(C)** the homogenized (diffuse) PL model described in the main text. **(D)** nSCLA vs IOP for the cases shown above. The top row **(A)** shows that, in the absence of a PL, SC collapse is predicted even at low values of IOP (< 15 mmHg); such collapse is not observed experimentally. The second row **(B)** shows that a single anteriorly-located PL is also not sufficient to prevent collapse at low IOP, with the two cases shown in **(A)** and **(B)** resulting in almost identical nSCLA responses vs. IOP (see **(D)**, noting the overlap between the blue solid line [No PL] and the orange dashed line [Single PL]). Only the diffuse (distributed) model of the PL ligaments avoided SC collapse at low IOPs (**C and D**). For panels **A-C**, colors represent separate tissues (e.g., PL is blue, see Figure 3).

**Figure S3:**
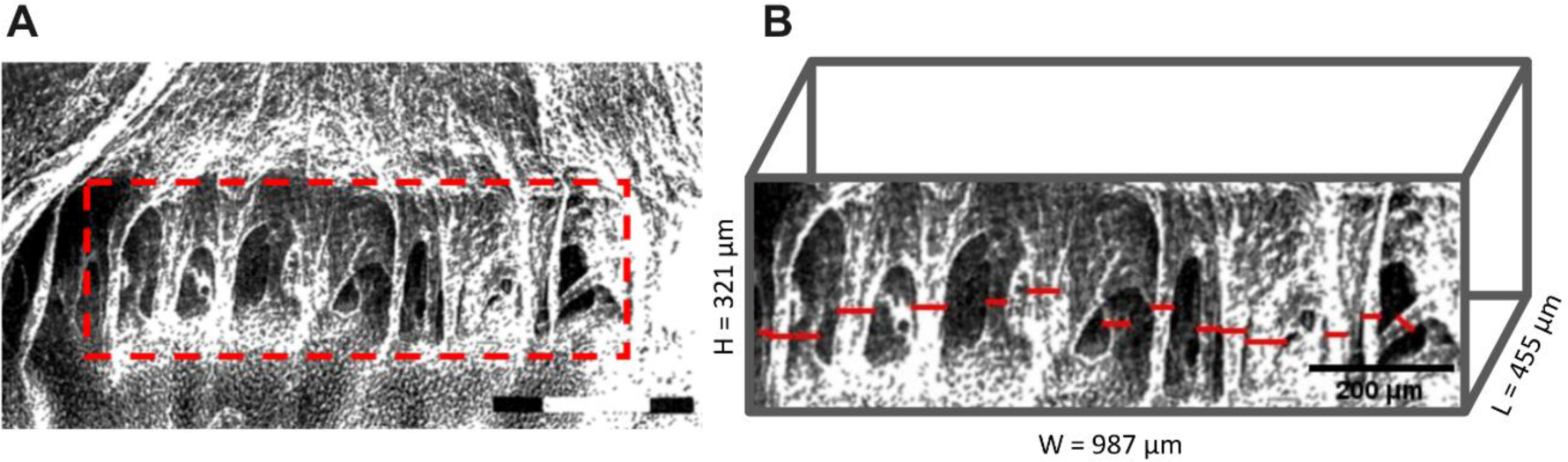
Description of the technique used to estimate PL solid fraction, β. **(A)** A Scanning electron micrograph of rabbit PL [1] with the PLs outlined by a dashed rectangle (red). White scale bar = 200 µm. **(B)** The radii of the PLs were determined by manually drawing line segments (red) in ImageJ, and the mean PL beam radius (*r*_*avg*_) was computed. The solid fraction within the relevant region of interest (ROI) was then determined by: (i) Projecting the red rectangle in panel (A) into the plane of the page by a distance *L*, determined from a histological image of the New Zealand white rabbit angle [2], showing the distance from the iridocorneal angle to the most anterior PL as *L* = 455 μ*m*. (ii) Computing β = ∑ *V*_PL_ /*V*_ROI_, where *N* is the number of PLs annotated in panel B (*N* = 15); ∑ *V*_PL_ = *N* × π*r*_*avg*_^2^*H* is the total volume of the PLs within the ROI, assuming cylindrically-shaped PLs; and *V*_ROI_ = *H* × *W* × *L* is the volume of the ROI. This procedure yielded β = 2.3%, which is clearly only an approximate estimate and should be refined in future work.

## Notes

### Competing Interest Statement

The authors have declared no competing interest.

### Summary of Updates

Revisions based on peer reviewed comments

